# Microvalve-based devices for flow-free gradient generators

**DOI:** 10.1101/2024.12.03.626578

**Authors:** Pierre Bohec, Florian Dupuy, Victoria Tishkova, Valentine Seveau de Noray, Marie-Pierre Valignat, Olivier Theodoly

## Abstract

Experiments with gradients of soluble bioactive species have significantly advanced with microfluidic developments that enable cell observation and stringent control of environmental conditions. While some methodologies rely on flow to establish gradients, other opt for flow-free conditions, which is particularly beneficial for studying non-adherent and/or shear-sensitive cells. In flow-free devices, bioactive species diffuse either through resistive microchannels in “microchannel-based” devices, a porous membrane in “membrane-based” devices, or a hydrogel in “gel-based” devices. However, despite significant advancements over traditional methods such as “Boyden chambers”, these technologies have not widely disseminated in biological laboratories, arguably due to entrenched practices and the intricate skills required for conducting microfluidic assays. Here, we integrated Quake-type pneumatic microvalves in place of microgrooves, membranes, or gels, and developed devices with precise control over residual flow, establishment initial gradient, and long-term stability of gradients. The “Microvalve-based” approach enables the generation of the automatization of delicate microfluidic manipulations, which paves the way for routine applications of controlled and tunable flow-free gradients in academic laboratories and biomedical units.

## Introduction

The ability of soluble biochemical gradients to influence the growth, motion and differentiation of individual cells or multicellular assemblies is crucial in most life-sustaining processes, such as foraging, immune recognition, wound healing or development. However, to study these phenomena in vitro, there is as yet no standard and user-friendly method for establishing controlled gradients at various temporal and spatial scales. The “Boyden chambers”^1^ that are the first in vitro assays targeting chemotaxis, consist of counting the fraction of cells crossing a microporous membrane in response to a solute gradient. Such end-point information without live imaging does not allow one to discriminate between chemokinesis (random migration) and chemotaxis (directed migration), and yields no precise information on migration parameters nor instant gradient shape and dynamics. Nevertheless, Boyden chambers have become a sort of standard method in the field and they are still widely used in biology laboratories. The “Zigmond chamber”^2^ was developed on the basis of the “Boyden chamber” to observe cell migration in real time, but it generates a short-duration gradient and is sensitive to evaporation. Later, the “Dunn chamber”^3^, the “Insall chamber”^4^ or the IBIDI’s “µ-slide chemotaxis” ^5^ were developed to enable long-term observation of cell migration and solve evaporation problems. Commercialized versions of these assays have found their way into academic biology laboratories, an important asset being that they require no special skills in micromanipulation. Still, time-zero conditions are poorly controlled, the gradient cannot be tuned once solutions are loaded, and/or cells may be crushed and intracellular factors released during assembly.

Microfluidic generation of chemical gradients was introduced around 20 years back^6^ and many devices^7^ have been developed to study the chemotaxis of various cells such as bacteria, neutrophils, T lymphocytes, dendritic, endothelial and cancer cells ^8–13^. A priori, the versatility of microfluidics is suited to design user-friendly and high-throughput devices for precise chemical gradients generation with long-term stability. However, there is no standard microfluidic device yet that satisfactorily combines all these features, and none are widely used nor commercialized. Existing microfluidic gradient generators can be divided into two groups: those that maintain stable gradients based on flow and diffusion, and those that establish local gradients using diffusion under flow-free conditions. The main advantages of flow-based devices ^14,15^ are their ability to generate gradients that are tunable and stable in space and time, together with their relative simplicity of handling. An intrinsic disadvantage of flow-based gradients is that they are not suitable for non-adherent or weakly adherent cells that can be expelled from the channels^16,17^, nor for mechanosensitive cells whose behavior may be altered by shear flow^18–21^, nor for long-term experiments as reagent consumption may become detrimental.

In flow-free devices, a test chamber designed to evaluate cell properties is usually separated from chambers containing bioactive species by pseudo-barriers that slow down flow (i.e. convection) while allowing diffusion^5^. The pseudo-barriers may be microchannels in “micro-groove-based” devices ^12,22,23,23–26^, a porous membrane in “membrane-based” devices^8,27–29^, or a hydrogel in “gel-based” devices^13,30–32^. A first problem with barrier-based devices is that the hydrodynamic resistance and physical extension of the barriers tend to flatten the gradients in the test chamber. This is most critical for microgrooves, which have both a high resistance and a large spatial extension, and less for membranes, which are thin, or for hydrogels, which are highly diluted and allow fast diffusion. A second challenge for barrier-based devices is to control the no-flow conditions strictly enough to respect the shape and stability of diffusion-based gradients because transport is allowed across them. Since pseudo-barrier cannot totally suppress convection, increasing their hydrodynamic resistance is a way to reduce it at will, but then the flattening of gradients is increased. Barrier-based devices require therefore to compromise between convection and diffusion control.

An efficient passive solution to cancel convection problems is to use dead-end microchambers^27^. In turn, this solution requires the device to be assembled in two stages, by loading cells into a first part of the device comprising the dead-end chambers, then implementing a second part of the device with the fluidic systems including barriers and bioreagents. A more flexible solution is to use microvalves to close the entrance and exit of the channels of interest, transforming channels into dead-end chambers^33^. This solution was successfully applied to a “gel-based” device to reduce residual convection^34^. In the present work, we went a step further and used microvalves not only to close inlet-outlet of channels (and transform channels into dead-end chambers), but also to replace pseudo-barriers separating the test chamber containing cellular specimen from the chambers with bioreactants. The former barrier-microvalves can be open to let the gradient build up by diffusion in absence of real physical barrier, the latter inlet/outlet-microvalves being locked to ensure dead-end chamber conditions. In turn, the barrier-microvalves can be closed to replenish the side chambers of bioreagents without perturbation of the gradient in the test chamber. This fully “microvalve-based” approach solves both the problems of residual convection and of hampered diffusion by physical barriers. Frank et Tay^33^ introduced a multiplexed system with barrier-microvalves and could establish 30 different conditions of gradients in parallel chambers. This important innovation was not followed by further dissemination and developments and applications, arguably due to handling complexity. Here, we developed “microvalve-based” flow-free generators that require no microfluidics skills from the experimentalists to allow a routine use for research or medical applications. Our devices use the standard design of a central channel as a test chamber with gradient, surrounded by two lateral reservoirs acting as chemokine source and sink ^5^. All delicate steps of fluid handling in microfluidic assays could be automated with microvalves. The single test chamber had a large size to accommodate thousands of cells, which is important for statistical strength in chemotaxis, and microvalves were automatically controlled to repeat experiments, screen different experimental conditions, and extend lifetime of gradients by refills cycles of biochemical channels.

## Results

### Rationale of microvalve-based design for gradient generation

Our devices are built on the generic design of a central test chamber surrounded by two lateral channels acting as bioreagent sources and/or sinks^5,34^ (Figure 1-A). Six inlet/oulet-microvalves are implemented to open/close the inlets and outlets of the three main channels, while two barrier-microvalves allow one to connect or disconnect the test chamber from the two lateral channels (Figure 1-A). All microvalves in this work are based on the Quake-Studer design, in which a PDMS membrane is pushed by pressure towards a fluidic channel with a rounded ceiling to promote tight closure^35^ (Figure 1-B). In one design, the microvalve-based barriers between the test chamber and the lateral channels are composed of a series of standard Quake-Studer microvalves with two actuation channels underneath the fluidic circuitry (Figure 1-C). The mold of the device is standardly made of SU-8 resin, except for the portion of channels in contact with microvalves that are made of AZT-40 resin to achieve rounded ceiling. In an alternative design, the test chamber and lateral channels are merged into a single chamber, and two actuation channels underneath separate this chamber into a central test chamber surrounded by two lateral channels under pressure (Figure 1-D). This latter device is simpler to fabricate, as the mold consists of a single layer of resin AZT-40 resin (Figure 1-B). In the two designs, connections for fluidic channels and microvalve actuations are similar, and the test chamber can be (dis)connected independently from the source or the sink channels. A picture of a full device filled with colored solutions is displayed in Figure 1-E (and Suppl. Mat. Movie 1 and Movie 2). Three pressure inputs allow one to open or close the inlet and outlet of the central, the source or the testing channels, whereas two pressure inputs can be operated to connect or disconnect the test chamber from the two side channels. In all, there are 11 tubing connections, 5 for microvalves actuation and 6 for fluidics control. Fluorescence microscopy was used to identify the suitable pressure range to operate microvalves (Figure 1F).

**Figure 1.**
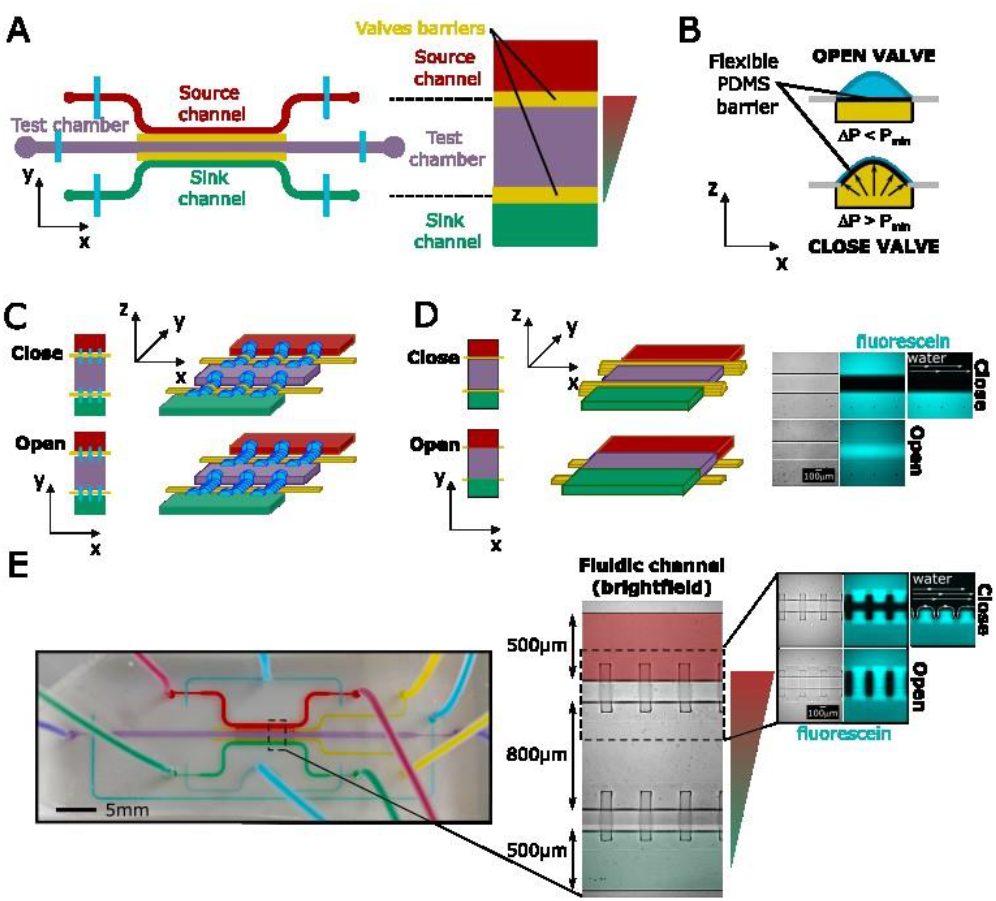
Device rationale of microvalves-based devices for gradient generation. **A**: Diagram of the microfluidics device depicting the test chamber in purple, the source channel in red, the sink channel in green, and the location of barriers in yellow. The inlets and outlets of the channels can be closed with inlet/outlet-microvalves (blue), while the test chamber can be (dis)-connected (from)-to the lateral channel by barrier-microvalves (yellow). B: The Quake-Studer pneumatic microvalves consist of a bilayer PDMS device with a flexible PDMS membrane (10µm thickness) separating the layers. Increasing the pressure in the bottom channel (yellow) causes the flexible membrane to clog the upper channel (blue). The upper channel has a rounded ceiling to facilitate the obstruction process. C: Left: 2D and 3D diagrams of device with test and sink/source channels connected with perpendicular rounded channels (blue), capable of being blocked by a bottom channel (yellow). D: Left-2D and 3D diagram of device with merged test and sink/source channels and barriers made by inflatable membranes underneath and across this chamber to form barriers. Right: Bright field and fluorescence imaging of device in non-inflated (open) and inflated (close) state (scale bar 100 µm). Once inflated, the adjacent channel formed can be washed with water without disturbing the other one. E: Left - Picture of the microfluidics device of figure C filled with colored solutions (scale bar 5 mm) showing the test chamber colored (purple), the source channel (red), the sink channel (green), the channels to actuate the inlet/oulet microvalves (blue) and the barrier-microvalves (yellow). Right - Brightfield image of the fluidic channel with corresponding dimensions. Top right - same as figure D for the design from figure C. (See also Suppl. Mat. Movie 1 and Movie 2)

### Refill of source/sink to maintain gradient boundary conditions

To obtain stable linear gradient by diffusion between a source and a sink, one needs to cancel transport (residual flows) and maintain stable boundary conditions (constant concentrations in source and sink). In practice, the concentration in the source and sink channels change due to diffusion, with a loss and a gain of material in the source or sink channel respectively. To restore initial conditions, we therefore carried out regular refills of the source and sink channels with fresh solutions via repetition of a two-step sequence of diffusion and refill (Figure 2A). During the “diffusion step”, all inlet/outlet microvalves were closed to cancel residual flow in the device, whereas the barrier microvalves connecting the test chamber and lateral channels were open to allow free diffusion from the source to the sink. During the “refill step”, the central channel was isolated from the rest of the device by closing the inlet/outlet microvalves of the test chamber and the barrier microvalves. A rapid refill (<10s) of the source and sink channels was possible without perturbation of the gradient in the test chamber. Refill/diffusion cycles were automated with a MATLAB interface controlling pressures transducer to actuate the microvalves, a Fluigent pumping system to inject solutions and a microscope to acquire images.

**Figure 2.**
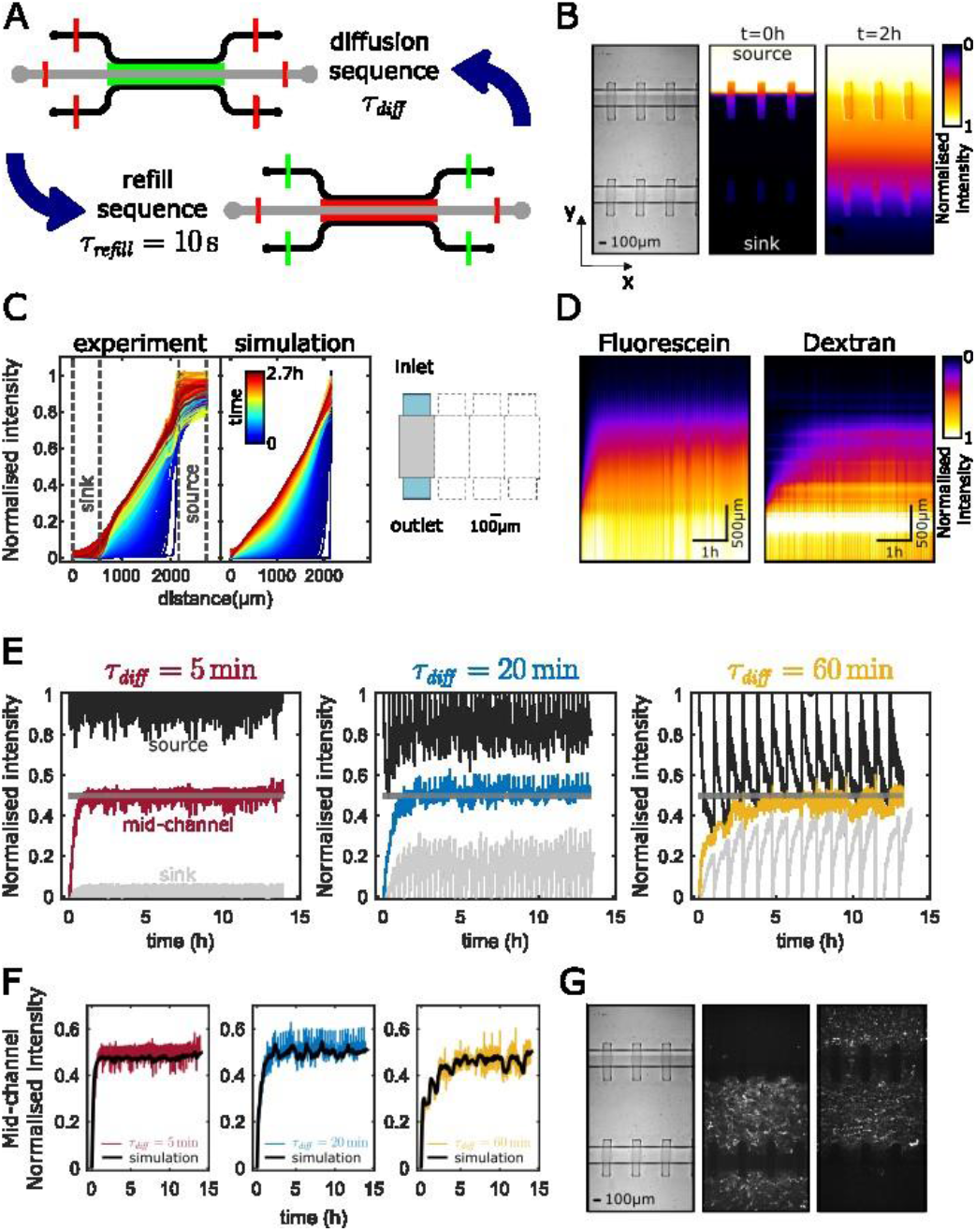
Gradient establishment kinetic and stability. A: Sequence diagram for maintaining a stable gradient. Diffusion sequence: The inlets/outlets microvalves of the source, sink, and test channelsar e closed (red), whereas the barrier-microvalves between the source/sink and the test chamber are open (green); Refill sequence: The inlet/outlet microvalves of the test chamber channel and the barrier microvalves are closed (red), whereas the inlet/outlet microvalves of the opened to enable the renewal of the liquids. With an inlet pressure of 200mbar, it took less than τ_refill_=10 seconds to fully refill the reservoirs. B: Left - Bright field image of the channels (scale bar 100µm). Middle – Fluorescence intensity of fluorescein in the channels at the beginning of the experiment. Initially (at t=0h), the source is filled with fluorescein in water (or the studied protein in cell media), the test chamber with water (or cells in media), and the sink with water (or cell media). Right – Fluorescence intensity of fluorescein in the channels after 2h of the gradient experiment (with a diffusion sequence duration τ_diff_=5min). The fluorescence intensity images are normalized by the average intensity of the source at t=0h. (See also Suppl. Mat Movie 3) C: Left - Normalized intensity profile measured with the fluorescein averaged along the x-axis as a function of the y-axis channel for different time points during the gradient experiment. Right – Box model used for the simulation of gradient concentration profile calculated using time-dependent finite element modelling (Comsol Multiphysics software), with central channel part (grey), source/sink channels part (blue), and dotted boxes to materialize the whole device as in Figure 2-B. D: Kymographs across the y-axis channel for 10kDa Fluorescein and 40kDa Dextran. The fluorescence intensity images is normalized by the average intensity of the source at t=0h. E: Normalized intensity with fluorescein at the middle of the channel over time averaged along the x-axis and average intensity in the source (black) andisink channel (grey) for different diffusion sequence durations : τ_diff_=5min (red), τ_diff_=20min (blue) and τ_diff_=60min (yellow). The intensities are normalised by the value of the source after each renewal. F: Same as E with the comsol simulation (black). G: Left - Bright field image showing the channels. Middle - Maximum projection of a 1-hour movie of 300nm GFP particles. Right - Maximum projection of a 5-minute movie showing the repeated sequence of opening and closing microvalves 8 times.

### Control of time-zero conditions

Before the start of an experiment, the barrier-microvalves between the test and lateral chambers were closed, the test chamber was loaded with cells, and the source and sink channels with the solutions of interest (Figure 2 B,C and Suppl. Mat Movie 3). The experiment started with a “diffusion step”. The initial temporal condition for the establishment of a gradient was therefore precisely defined by the instant of barrier-microvalve opening, while the initial spatial conditions where precisely controlled as a step profile positioned on the axis of symmetry of the barrier-microvalves. Precise control of spatio-temporal time-zero conditions is an important asset to probe rapid cellular responses to biochemical cues or drugs.

### Transient durations can be tuned by the refill/diffusion sequences

The time to reach steady state conditions (linear gradient between the source and sink) depends on the geometry of the device and the diffusion constant of the species of interest, as exemplified with fluorescein and a fluorescent 10kDa dextran in Figure 2-D. However, our devices also enabled adjustment of the transient period duration by modifying the refill/diffusion sequence. Figure 2-E reports the kinetic of gradient establishment with the same 10 kDa dextran and three different sequences consisting a similar refill step of 10s and diffusion steps is of 5, 10 or 20 min. As expected, the depletion in the source channel and the accumulation in the sink channel during the diffusion sequences increased significantly with the duration of the diffusion sequence. Hence, as the interval between refills was increased, the experiments deviated further from the ideal scenario of stable boundary conditions. Yet, the concentration in the test chamber reached a value very close to 0.5 at steady state, meaning that the final state remained very close to the ideal case of linear gradient between a source and a sink of constant concentrations. This demonstrates the possibility of tuning the duration of the transient state while reaching similar steady state conditions, which may be useful to screen a wider range of gradient conditions in a single experiment or to test phenotypes at different gradient loading rates.

### Gradients are diffusion-controlled and stable on the long term

The gradient establishment in the devices was compared to simulations with COMSOL using geometrical elements of Figure 2-C. The transient states match well between experiments and simulations (Figure 2-C,F), which validates the fact that the flow-free conditions were strict enough to ensure that gradient establishment was dominantly controlled by diffusion. Furthermore, the equilibrium state of the gradient remained remarkably stable over the whole experiments (that was stopped here after 14 hours).

### Flow-free conditions allow probing non-adherent cells

Flow-free conditions can typically be perturbed by local inhomogeneities of temperature, pressure and concentration, as well as by water evaporation through the PDMS^36^. To assess flow during the refill and diffusion steps, the devices were loaded with a solution of 200 nm diameter fluorescent beads. With inlet/outlet microvalves closed and the use of an OKOLAB chamber to maintain stable temperature and humidity conditions, flow drifts were lower than 0.1 µm/min (Figure 2-G) during diffusion or refill steps, which was low enough to record slow processes such as leukocytes swimming (speed of 5µm/min). However, the switch between refill and diffusion steps induced fluid displacements due to the motion of the microvalve membrane (Figure 2-G). Beads or cells in the test chamber moved typically by 15 µm +/-5µm in one direction upon barrier-microvalve closing at the start of the refill step, then moved in the other direction at the end of the refill step, with a resulting displacement per cycle is of 19 µm +/-6 µm. This perturbation has no impact on experiments that consider only the instant properties of cells and not the continuous tracking of individuals cells during several refill/diffusion sequences. In turn, when cell tracking on several refill/diffusion cycles is required, the perturbation can be minimized by lowering the frequency of refill/diffusion cycles, or readily corrected during data process by using the information of microbeads tracking as corrective reference.

### Experiment of chemokinesis and chemotaxis

To test the functioning of the devices, we performed chemotaxis assays with non-adherent primary human neutrophils swimming in a gradient of FMLP (Figure 3 and Suppl. Mat. Movie 4 and Movie 5). The spatiotemporal properties of the FMLP gradient were determined experimentally using fluorescein in the source as fluorescent marker of diffusion. This enabled to assess the local average concentration and the local gradient of concentration in the whole test chamber at each instant of the experiment (Figure 3A). Cells were individually tracked to determine their instant orientation and velocity (Figure 3B), while cells shapes allowed us to identify their polarization state, a round shape corresponding to quiescent state and polarized shape to motile^37,38^ (Figure 3B). To assess their orientation versus a chemical gradient (here along the y-axis), we calculated a chemical index, CI, defined as the distance traveled along the chemokine axis (Δy) divided by the total trajectory length (Figure 3D). At least 1000 cells were tracked per experiment thanks to the large size of the test chambers, and in order to further cumulate strong statistics the experiments were run several times by automatic refilling of test chambers with fresh cells. In the initial moments, the arrival of FMLP triggered the polarization and motility of cells. This which is characteristic of a chemokinesis effect and consistent with data in homogenous concentration (Figure 3-C). Interestingly, the average velocity of polarized cells themselves was enhanced with FMLP concentration, which suggests that chemokine sensing can tune different levels of polarization and motility. Besides, the cells were guided towards increasing concentration of FMLP, which is characteristic of a chemotactic effect of FMLP (Figure 3-E). The average guiding effect increased with concentration in the range 0-100 nM. We built 2D plots of chemical index versus the local average concentration and the local gradient steepness surrounding cells (Figure 3-F). Experiments with different concentration in the source channels were performed to widen the range of gradient conditions. For the experiment with 1 nM FMLP in the source channel, the CI was null at lowest local concentration and lowest gradient steepness, which is expected because there can be no guidance when the cue is absent or too weak. It is then surprising at first that the CI were not zero in experiments with 10 nM FMLP at lowest gradient steepness, as well in experiments with 100nM FMLP at both lowest local concentration and lowest gradient steepness. In fact, the size of the boxes and the scales in the 3 plots of Figure 3G are very different, and the conditions of null CI are hidden due to pooling of other finite values in the same box plot. These remarks underline the importance of performing experiments at different concentration to explore cell phenotypes at relevant ranges and resolutions.

**Figure 3.**
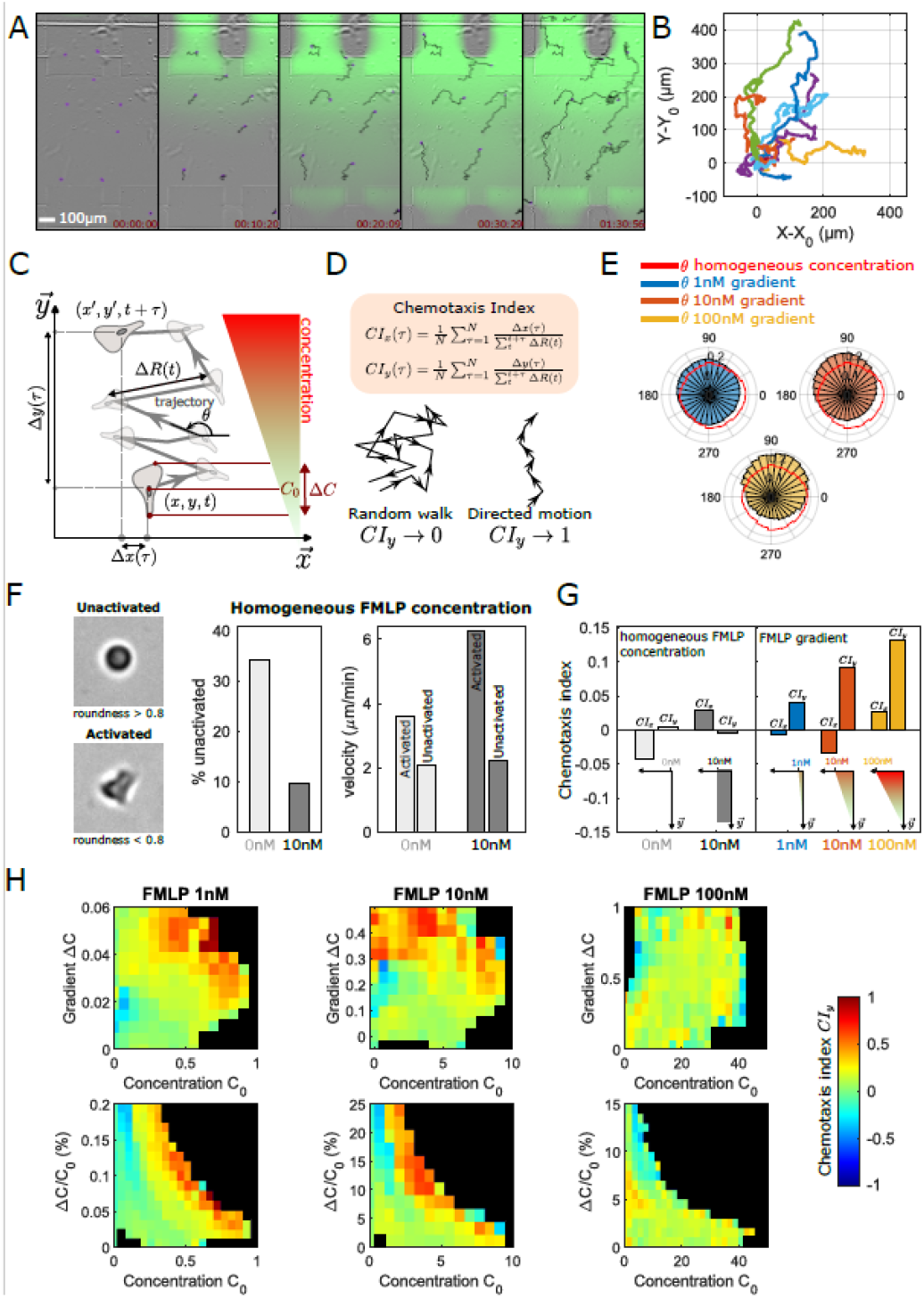
Neutrophil chemokinesis and chemotaxis. A: Time-lapse of bright-field images capturing neutrophil migration in an FMLP gradient with a source at 15 nM (scale bar 100µm). Trajectories are highlighted in black to indicate migration. Fluorescence intensity of fluorescein is superimposed on the image to visualize the FMLP gradient. (See also Movie 3 and Movie 4) B: Trajectories from the experiment in A, originating at the origin (n=9). C: Left - Images of activated (polarized) or unactivated (unpolarized) neutrophil cells differentiated by their roundness level (threshold of 0.8). Right - Percentage of unactivated cells and cell velocity (µm/min) for the unactivated and activated cells in a homogeneous profile of 0nM (pale grey) and 10nM (grey) FMLP concentration. D: Diagram explaining the trajectory analysis in a gradient. We define the displacement Δx(τ) Δy(τ) of the cell during the interval time τ and the local gradient ΔC. E: Definition and illustration of the chemotaxis along x-axis and y-axis index for the interval time ⍰. F: Angle distribution for an interval time ⍰=1min for gradient experiments with 1nM (blue, N_track_=22886, n=628981), 10nM (orange, N_track_=29300, n=624699), and 100nM (yellow, N_track_=2007, n=89659) FMLP at the source, and for experiments in a homogeneous concentration of 10nM FMLP (red, N_track_=1446, n=63142). G: Bar plot of the average of the chemotaxis index along the x-axis (CIx) and y-axis (CIy) in a homogeneous profile of 0nM (pale grey, N_track_= 1877, n= 130349) and 10nM (grey, N_track_=1446, n=63142) FMLP concentration, and in a gradient profile of 1nM (blue, N_trac_k=22886, n=628981), 10nM (orange, N_track_=29300, n=624699), and 100nM (yellow, N_track_=2007, n=89659). The chemotaxis indices are calculated for a time interval of 1min. H: Heatmaps of the chemotaxis index as a function of the local concentration and - Top-the local gradient difference ΔC in front of and behind the cell; or - Bottom- the local gradient steepness ΔC/C. Cases in black indicate missing values.

Please do not adjust margins

### DISCUSSION

The generation of flow-free gradients is attractive for applications with cellular samples that are non-adherent, shear sensitive, or influenced by a contact with a substrate. Yet, it is challenging to achieve satisfying flow-free conditions, especially for experiments with slow diffusing chemicals and slow swimming specimen. We have shown here that microvalve-based devices offer unique solutions to overcome gradient instability, flow drifts, uncontrolled boundary conditions, low statistics, and complex experimental handling.

Inlet/outlet microvalves were used in our devices to minimize convection. With a channel width L= 800µm, a residual fluid transport of u =0.1 µm/min and a typical diffusion coefficient D = 90 µm^2^/s, the typical Peclet number was equal to 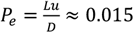, which corresponds diffusion-controlled regime. Establishment of protein gradients under diffusion control was attested by comparisons with COMSOL simulations of pure flow free conditions. Hence, microvalves-based devices combined the performance of dead-end chambers for flow-free conditions^27^ with the flexibility of achieving dead-end conditions on demand and with the possibility to reload the device to change/repeat experimental conditions. Satisfying flow-free conditions have previously been enabled by fully immersing devices in medium^13,34^, however such cumbersome manipulations are unnecessary with microvalve-based devices.

The barrier-microvalves (between test chamber et source/sink channels) form the second important asset of our device to optimize the spatio-temporal control of gradients establishment. At experiment onset, reservoirs of biochemicals are duly filled with stock solution while disconnected from the test chamber, then the opening of barrier-microvalves defines clear time-zero conditions with step concentration profiles between cell chamber and sources. This is a significant asset as compared to loading of devices via manual pipetting that involves poor control of initial convection and diffusion conditions. Our microvalves-devices are therefore particularly well-suited for studying rapid cellular responses to drugs, cytokines, or chemokines.

Importantly, microvalves-based devices are suited for automation. This was crucial to maintain long-term gradient stability by regular refill of sources and sinks. The advantage of refills as compared to the use of large reservoirs to delay the depletion of bioreactants^5,26^ is that there is no time limit associated to the size of the reservoirs. Long-term stability is interesting to study chemotaxis of mesenchymal cells, cell differentiation, or development of organoids and embryos. Automation is also an asset to repeat experiments and screen of experimental conditions (gradient characteristics, cells types and reactants), which is important to analyze noisy phenotypes such as chemotaxis^39^ and to screen parameters for combinatorial studies.

Microvalves devices were used here to analyze chemokinesis and chemotaxis of non-adherent primary neutrophils. The large test chamber hosting manyicells and automated repetition of experiments allowed a thorough inedite quantitative characterization of the noisy chemotaxis of swimming leukocytes. The polarization, speed and guidance of neutrophils versus FMLP were characterized for a wide range of concentrations (< 0.1nM to > 40 nm) and gradient steepness (ΔC/C between around 0.01 and 1). The optimum of chemotaxis was found at concentrations 1-5 nM and steepness ΔC/C between 0.05-0.5, which is consistent with previous data^38,39^, whereas the relatively lower CI values can be explained by the larger rotative diffusion of non-adherent cells as compared to adherent crawling cells in previous studies.Finally, the complexity inherent to microfluidic flow-free assays is managed here by thoughtful design and automation of the device. Critical aspects such as minute fluid handling, drift cancellation, control of initial and limit conditions, elimination of bubbles (a microfluidicist nightmare), and data analysis can be handled though pre-registered automatized routines. The tedious operation of plugging all tubing could be easily streamlined with the introduction of a specially engineered connector capable of simultaneously plugging all tubing. Microvalve-based flow-free gradient generators, with their assets of short- and long-term control and of high-throughput measurement are therefore suited to disseminate towards research and biomedical laboratories that are non-expert in microfluidics.

## Material and methods

### Fabrication of the PDMS microdevices

Microfluidic devices were produced by soft photolithography and micromolding techniques. Molds were fabricated by spin coating a layer of the negative photoresist SU-8 (MicroChem Newton, MA) (SU-8 3050, h=50 µm) on a 4-inch silicon wafer (Siltronix). The photoresist was exposed to UV light through the mask containing the gradient design. SU-8 developer solution (MicroChem, Newton, MA) was used to dissolve unexposed parts of the photoresist. To obtain channels with rounded edges for microvalves we spin-coated a positive photoresist AZ-40XT at 500 rpm for 10s and 2800 rpm for 20s. Spin coating was followed by a 7min baking at 125°C, alignment and exposure to UV for 30s, a second bake for 1min and development in AZ326 MIF for 3.5min followed by a third bake for 7min. Microvalve molds were produced on a separate wafer using SU-8 3050, h=40 µm. PDMS molding was performed by mixing the pre-polymer (Sylgard 184, Dow Corning) with the polymerization agent at 10:1 ratio for the devices and 12:1 ratio for the microvalve master molds. PDMS was directly poured on the device molds, followed by degassing in a vacuum bell and spin coated on the microvalve molds to obtain a thickness of 10 µm above the channels (actuator membrane). Molds where baked in a 65°C oven for at least 2 hours. After curing, only the fluidic devices were unmolded and both PDMS surfaces (devices and microvalve layers) were treated in UV-Ozone for 30min, overlaid on the microvalve mold and left overnight at 65ºC to assure strong bonding. The next day the PDMS montages were removed from the microvalve molds and the inlets and outlets were punched with 1.2 mm punchers (Harris Uni-Core), before being sealed on a glass slide via plasma activation (Harrick Plasma) for 15min and final 95ºC baking for another 15min.

### Surface treatments

In order to deter cell and protein adhesion for experiments with non-adherent cells, fluidic circuitry was filled with a 4% Bovine Serum Albumine solution for 20 min then rinsed with Milliq-water and filled with medium.

### Reactants

FMLP (Sigma Aldrich) solutions were prepared in culture media at 1, 10, and 100 nM. Fluorescein (376 Da, Sigma) solution at 0.1 mg/ml was added to monitor monitor gradient dynamics.

### Cells

Whole blood from healthy adult donors was obtained from the “Établissement Français du Sang”. Neutrophils were extracted with the EasySep™ Direct Human Isolation Kit (STEMCELL Technologies), following manufacturer instructions. After purification, cells were kept in RPMI 1640 medium supplemented with penicillin 100 U/ml (Gibco, Carlsbad, CA), streptomycin 100 µg/ml (Gibco, Carlsbad, CA), 25 mM GlutaMax (Gibco, Carlsbad, CA), and 10% fetal calf serum (FCS; Lonza, Basel, Switzerland) in a 37°C incubator with 5% CO2, until use.

### Flow control and microvalves operation

The channels for microvalve actuation were filled with water and controlled using electro-fluidic microvalves (The Lee company), a Velleman card (P8055N-2), and a pressure source of 1.5 bar. The fluidic circuitry was fully filled with medium and connected to a microfluidic pressure control system (Fluigent, MFCS-EZ). To ease the experiment procedure, a graphical user interface was developed using MATLAB (see Supplementary Figure 1), allowing for the opening and closing of all microvalves and the initiation of diffusion and refill sequences.

### Imaging and data analysis

Experiments were conducted using an inverted Zeiss Z1 automated microscope (Carl Zeiss, Germany) equipped with a CoolSnap HQ CCD camera (Photometrics) and controlled by µManager^1.4^. Bright-field and fluorescent images were acquired using Plan-Apochromat 10x/0.3 and 20x/0.8 objectives with images captured at 10-second intervals. RICM images were processed as follows: first, illumination correction was performed by subtracting a background image, and then the images were binarized using Labkit^42^, a FIJI plugin for Pixel classification. Cell tracking was conducted using the FIJI plugin Trackmate^40^.

Tracks were exported and further analysis and plots were conducted using a custom-made MATLAB script (MATLAB software, The MathWorks, Natick, MA, USA). Gradient dynamics were analysed with another custom-made MATLAB script. Fluorescence intensity values were extracted for each time point along a line drawn in the direction of chemokine diffusion, with a width of 1024 pixels to average camera noise. The intensity values were then normalized using the equation:

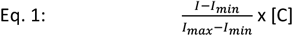

Where *I*_*min*_ is the average value recorded on the sink channel after each refill sequence (background), *I*_*max*_ is the average value recorded on the source channel after each refill sequence, and [C] is the chemokine concentration applied at the source.

### Modeling

Gradients of concentration were calculated using time-depended finite element modelling (Comsol 6.1 Multiphysic software)^5.4^. The 2D geometry was drown directly in Comsol 6.1 and is represented in Figure 2c by a colored part. The total length of the modeled geometry is 1500 µm, with a width of 300 µm for the inlet/outlet and 400 µm for the central part. The concentration gradient was calculated by solving Fick’s diffusion equation: ∇·(−*D* · ∇*c*) + *u* · ∇*c* = *R*, where *D* is the diffusion coefficient, *c* – concentration of the species and *R* is reaction rate which is set to zero in our mode. Convection was not considered, and fluid flow velocity *u* was set to zero. Free triangular mesh with 8530 elements with element sizes ranging from 12 to 17 µm. The minimum and maximum skewness are 0.64 and 0.94, respectively. A perfect case is referenced with skewness equal to 1. Time-dependent solver with strict time step of 1s was used for all the calculations. The MUltifrontal Massively Parallel sparse direct Solver (MUMPS) was used. Concentration on the inlet(source) and outlet (sink) boundary conditions were assigned using experimentally measured fluorescence on the opening and closing microvalves. The diffusion coefficient *D* was set to 480 µm^2^/s. The results of the concentration gradient have been normalized. The microvalve opening and closing cycles of 5 min, 20 min and 1 hour have been modelled for the total duration of 14 hours each.

## LIVE SUBJECT STATEMENT

Human subjects: Blood from healthy volunteers was obtained through a formalized agreement with French Blood Agency (Etablissement Français du Sang, agreement n° 2017-7222). Blood was obtained by the agency after informed consent of the donors, in accordance with the Declaration of Helsinki. All experiments were approved by the INSERM Institutional Review Board and ethics committee.

## Author contributions

PB and OT designed and optimized the devices, PB and FD fabricated the devices, PB wrote all the scripts to automatize the devices, PB and FD performed the experiments, VSdN helped with microfluidic experiments, VT designed and performed COMSOL simulations, MPV contributed to project design and supervision, OT wrote the manuscript and supervised the project.

## Acknowledgements

This work was supported by Agence Nationale de la Recherche (grants RECRUTE AAP CE15); LABEX INFORM; Région PACA and the Turing Centre for Living systems. Also, it has been carried out with the financial support of the Regional Council of Provence-Alpes-Côte d’Azur and with the financial support of the A*MIDEX (n° ANR-11-IDEX-0001-02), funded by the Investissements d’Avenir project funded by the French Government, managed by the French National Research Agency (ANR).. We also thank Frederic Bedu, Igor Ozerov and Gérald Clisson for welcoming us at the microfabrication facilities of LOF (CNRS-Solvay, Bordeaux) and Planete Cinam (CNRS-AMU, Marseille), and Thi Tien N’Guyen for welcoming us at the Cell Culture Platform facility (PCC, Luminy TPR2-INSERM, Marseille).

## Supplementary movies

Movie 1: **Functioning of microvalves-based devices for gradient generation-general view**. The device corresponds to design of Figure 1 C and picture of Figure 1E. The channels are filled with colored to distinguish the test chamber (white), the source channel (red), the sink channel (green), the channels to actuate the inlet/oulet microvalves (blue) and the barrier-microvalves (yellow). At time zero, all inlet/oulet microvalves (blue) and the barrier-microvalves (yellow) are closed and the central channel is not colored. At time XX, the barrier valves (yellow) are opened and the red and green dye start to diffuse in the central channel. At time XX, barrier valves (yellow) and Input/output valves of the sink and source are closed, then the input/output valves of the central channel are opened to rinse the channel with purple fluid. Valves appear blue or yellow when closed for inlet/outlet channels and barrier channels respectively, whereas the color of the fluidic channel appears when they are open.

Movie 2: **Functioning of microvalves-based devices for gradient generation-general view – Zoom on central chamber**. This movie corresponds to the same data as in Movie 1 with a close-up of the test chamber. One can see that at time zero the barrier-microvalves (yellow) are closed, then at time XX, they are opened allowing diffusion from the side channel (red and green), and at time XX, they are closed again and the central channel is filled with purple fluid. Barrier microvalves appear yellow when closed, whereas the color of the fluidic channel appears when they are open.

Movie 3: **Establishment and maintenance of a fluorescein gradient with cycles of diffusion and refill steps - Quantification**. The data are taken in a device corresponding to Figure 1-D, with barriers consisting of single long micro-valves for the sink and the source. Left-Videomicroscopy in epifluorescence mode focusing on the central part of the device comprising the central channels, the sink and source microvalves and part of sink and source channels. The sequence starts with a central channel free of fluorescein and shows the establishment and maintenance of a fluorescein gradient between a source at 15 nM (Bottom) and the sink at 0 nM (top). At each refill step, the microvalve appear dark when closed since channels bottom and floor are in contact and fluroscent fluid is expelled. Time is in hrs:min:s. Right: timelapse of fluorescence intensity profile taken along the vertical yellow line superimposed on the videomicroscopy data.

Movie 4: **Neutrophil chemokinesis and chemotaxis – focus on a few cells**. Time-lapse of bright-field images capturing neutrophil migration in an FMLP gradient with a source at 15 nM (scale bar 100µm). Trajectories are highlighted in black to indicate migration. Fluorescence intensity of fluorescein is superimposed on the image to visualize the FMLP gradient.

Movie 5: **Neutrophil chemokinesis and chemotaxis – typical experiments with large populations**. Time-lapse of bright-field images capturing neutrophil migration in an FMLP gradient with a source at 15 nM (scale bar 100µm). Trajectories are highlighted in black to indicate migration. Fluorescence intensity of fluorescein is superimposed on the image to visualize the FMLP gradient.

